# Explainable machine learning reveals an RBP regulatory logic of exon skipping

**DOI:** 10.64898/2026.05.29.728731

**Authors:** Yogindra Raghav, Ayan Paul, Reece Anderson, Shalini Karthyk, Annette Iturralde, Janmejay Vyas, Jennifer Dy, Brett C. Jones, Peter J. Castaldi, John Platig

## Abstract

RNA binding proteins (RBPs) regulate the life cycle of an mRNA, often through RBP-RNA interactions. This life cycle includes splicing, whereby the intronic sequence of a pre-mRNA is removed and the exons are joined together. However, the patterns of RBP binding that lead to different splicing outcomes are still incompletely understood. Here, we build machine learning models from RBP-RNA binding and knockdown RNA-seq data for over 168 RBPs in two cell lines (HepG2 and K562) to better understand the binding patterns that predict exon skipping, the predominant form of alternative splicing in humans. We show that models trained exclusively on RBP binding patterns are indeed predictive and that a more sophisticated machine learning model (XGBoost) outperforms simpler linear models. In addition, we are able to extract a biologically interpretable logic embedded in these models. We show that SHAP, a machine learning explainability technique, captures activating and repressive behavior of RBP binding that is position-specific. In addition, we find that SHAP values are predictive of changes in unseen splicing events and that SHAP interactions between pairs of RBPs are predictive of protein-protein interactions. Our results demonstrate that using machine learning with interpretability techniques can reveal a regulatory logic of RBP binding. By estimating the impact of an RBP binding site on a splicing event, the SHAP values also provide a directly testable scientific hypothesis. We anticipate that models designed around biological processes and focused on interpretability will yield actionable biological insights both in splicing and genomics generally.

## 1 Introduction

RNA binding proteins (RBPs) are critical regulators of an mRNA’s life cycle. This includes RNA splicing, where intronic sequences are removed from pre-mRNAs through recruitment of the spliceosome. This process also enables cells to alternatively splice transcripts by selectively removing exons, allowing an individual gene to generate a diversity of transcripts. Estimates based on RNA-seq data suggest that ∼95% of multi-exonic genes are alternatively spliced, resulting in ∼100,000 alternative splicing events that contribute to proteome diversity in major human tissues [1]. Roughly half of all expressed genes have tissue-specific isoforms [2], and alternative splicing has been shown to play an important role in cancer and other diseases [3–5]. In addition, disease associated variants are linked to changes in splicing [3, 6]. There is also interest in identifying pathological alternative splicing events that can be targeted using antisense oligonucleotides (ASOs) [7, 8].

Given the essential nature of alternative splicing and its therapeutic potential, there has been a longstanding interest in using the nucleotide sequence around splice sites to predict instances of alternative splicing [9–14]. These approaches have made limited use of RBP binding, largely due to a lack of available data. However, recent efforts to widely profile the alternative splicing effects upon RBP knockdown [15, 16] and RBP binding [17] have enabled the direct modeling of the impact of RBP binding on alternative splicing. This includes models predicting the impact of using RBP (eCLIP) binding from cancer cell lines to predict tissue specific changes in brain cell types [18] and integration of RBP knockdown data with RBP sequence motifs to explain the splicing changes that occur after a perturbation experiment [19].

Motivated by this, we aim to answer two questions: (1) To what extent can we predict exon skipping using RBP binding patterns derived from cell-line-matched eCLIP? (2) Can we extract a meaningful RBP regulatory logic from these prediction models using explainable machine learning (xAI) techniques? To address the first question, we tested a tree-based machine learning model on its ability to predict exon skipping from the pattern of RBP binding surrounding the skipping event (see Figure 1). Given recent examples in other genomics contexts [20] of linear models outperforming more complex machine learning models, we first compared our model to simpler linear models.

**Figure 1.**
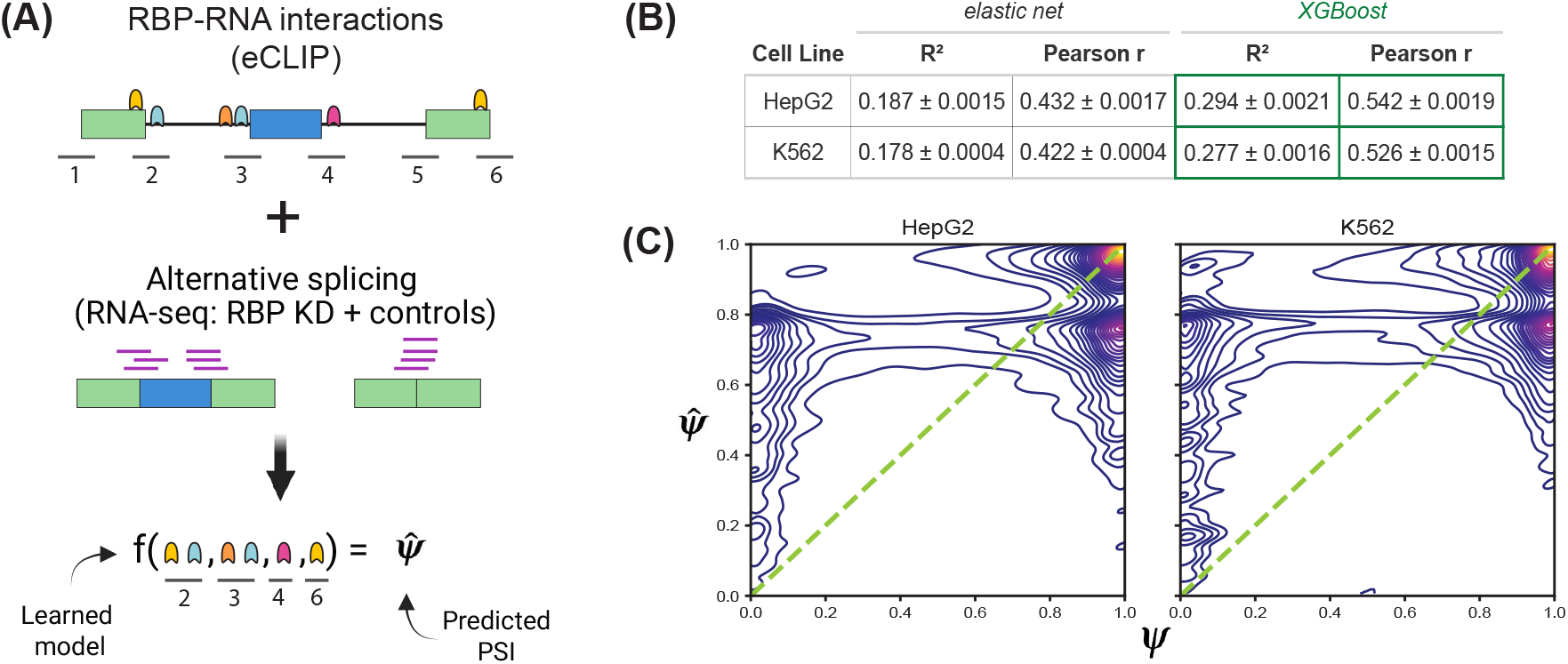
Overview of modeling approach and performance. **(A)** Illustration of modeling approach. ENCODE-derived eCLIP [17] binding sites were mapped to windows ±100bp around exon boundaries (“positions”, labeled 1-6) for skipped exon (SE) events measured in shRNA RBP knockdown and control RNA-seq samples. Models were then trained to predict the Percent Spliced In (*ψ*) for an SE event from an RNA-seq sample using only the corresponding RBP binding pattern. If an RNA-seq sample measured an RBP knockdown, the binding sites for that RBP were set to zero (Methods 4.2). **(B)** Average performance of the top 5 models on held out (test) data (Methods 4.3.1). **(C)** Scatterplots of actual (*ψ*) and predicted 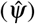 values within the “test” dataset for the best model per cell line.

While other models have incorporated RBP binding derived from limited CLIP-seq [21] as part of larger models, in this work, our aim is to explore the extent to which positional RBP binding information alone can predict exon skipping. The advantage of this data-restricted approach is that it directly interrogates the value of large-scale RBP binding profiles and simplifies the interpretability of the models learned from these data.

To address the second question of identifying a regulatory logic for RBPs, we used SHapley Additive exPlanations (SHAP) [22] to identify binding sites that contributed to the predictions from the model. We also assessed the functional relevance of these binding sites on splicing events from unseen RBP knockdowns. Using the SHAP values, we then looked for master splicing regulators and position-specific RBP effects (e.g., RBP binding that mattered at some splice sites but not others). Finally, we used SHAP to calculate RBP-RBP interactions and look for position-specific, co-operative effects of RBP-RBP regulation. We then compared these interactions with known protein-protein interactions.

## 2 Results

To model exon skipping as a function of position-specific RBP binding, we retrieved eCLIP and shRNA knockdown (KD) RNA-seq data from ENCODE [15]. In HepG2, we downloaded eCLIP data for 105 RBPs and knockdown RNA-seq (KD and control samples) for 85 RBPs. In K562, we downloaded 139 eCLIP and 101 shRNA data sets. All RBPs with knockdown RNA-seq were a subset of the eCLIP RBPs (Methods 4.1). Percent Spliced In (*ψ*) values for the knockdown RNA-seq data were calculated using rMATS [23] (Methods 4.1.1).

For each Skipped Exon (SE) event observed in an RNA-seq sample, we labeled 200 base pair windows centered on the exon boundaries of each of the three exons involved in the skipping event to get 6 “positions” (Figure 1A) and then used eCLIP data to indicate which RBPs were bound at each of the six positions (Methods 4.2). In total, we compiled 4.57 million (HepG2) and 6.29 million (K562) instances of an RBP binding pattern paired with a *ψ* value from an SE event. We define a “Splicing Instance” (SI) to be a binding pattern around an SE event paired with the SE event’s condition-specific *ψ* value. For an SI where the *ψ* value came from an RBP knockdown RNA-seq sample, we set the binding value of the corresponding RBP to zero (“in silico knockdown”) at all positions (Methods 4.2). The quantiles for the number of binding events per SI are shown in Figure S1A. For our downstream work, our models took as input an SI’s position-specific RBP binding pattern to predict the corresponding *ψ* value. This approach tests the extent to which SE splicing can be predicted by using RBP binding to specific locations along an SE event and provides the ability to tease out novel regulatory logic around the position-specific effects of RBPs on their own and in conjunction with other RBPs.

### 2.1 Tree-based models outperform linear models in predicting *ψ* from RBP binding

Before applying more sophisticated machine learning models, we first tested the performance of using a penalized linear model (elastic net) where each model input (feature) is the binding of a given RBP (e.g., QKI) at a specific position (e.g., 5’ splice site of the upstream intron – position 2 in our notation; Methods 4.3.1). Our elastic net model then took either 630 (6 × 105) HepG2 or 834 (6 × 139) K562 features as inputs and was trained on 80% of autosomal genes and tested on the remaining 20% (“test” dataset; Methods 4.3). Throughout this work, we created separate data sets and models for each cell line. We found the performance of our top 5 elastic net models to be similar between cell lines with an *R*^2^ value around 0.18 (Figure 1B).

Next, we used a tree-based approach, XGBoost [24], again providing binarized binding data as an input and predicting *ψ*. After training and tuning these models (Methods 4.3.1), XGBoost significantly outperformed elastic net in both cell lines (Figure 1B). Like elastic net, the XGBoost models perform similarly between cell lines. Given its improved performance, we used XGBoost for all downstream analyses. As shown in Figure 1C, our models predict highly-included *ψ* splicing events well but also tend to generally predict values in the upper end of the *ψ* range, potentially due to the imbalanced distribution of *ψ* (Figure S1B).

### 2.2 Explainability (SHAP) scores can predict RBP-driven splicing changes

We next sought to interrogate which binding sites the XGBoost models used when making their predictions. To do this, we used the SHapley Additive ExPlanation (SHAP) method [22]. Similar to a term in a regression model, SHAP provides an estimate of how much a given feature (an RBP binding at a position) contributes to a specific prediction. In other words, given a binding pattern, how much does each feature contribute to the resulting *ψ* prediction? For further details regarding SHAP, please see Methods 4.4 and 5.1.

We hypothesized that the SHAP values derived from our XGBoost models could consistently identify “functional” RBP binding sites. Here, we define functional to mean an RBP’s binding site(s) within SE events that were differentially spliced (Δ*ψ*) upon knockdown of that RBP. To test this hypothesis, we took all SE events unseen by the model (test data) and obtained the corresponding SHAP values for the binding sites within those events. We then compared those SHAP values to the differential splicing (Δ*ψ*) outcome of those events upon knockdown of that RBP (Methods 4.5). When comparing the direction of effect (increasing/decreasing *ψ*) of the SHAP values to the sign of statistically significant (FDR ≤ 0.1) Δ*ψ*, we found positive correlations across a range of SHAP value cut-offs in both cell lines (Figure 2A).

**Figure 2.**
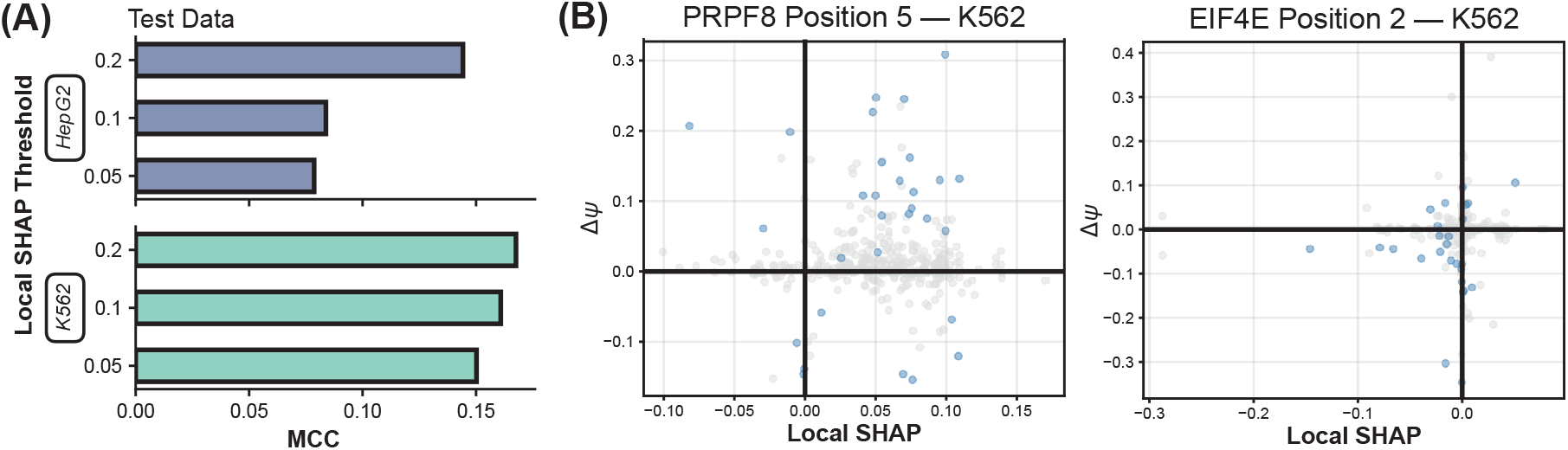
Local SHAP values at individual RBP binding sites are predictive of RBP perturbation effects. **(A)** Matthew’s correlation coefficient (MCC) comparing the sign of an RBP’s local SHAP value (at a given position) with the change in splicing (sign of Δ*ψ*) upon knockdown of that RBP (Methods 4.5). Comparisons are made using only events from “test” data and that showed statistically significant splicing changes (rMATS FDR value ≤ 0.1) upon knockdown. **(B)** Scatterplot of local SHAP vs Δ*ψ* for unseen knockdowns of PRPF8 and EIF4E using CRISPRi knockdowns (Methods 4.6). Significant events are labeled in blue (rMATS FDR value ≤ 0.1).

As a further test of the SHAP values, we downloaded CRISPRi RBP knockdown data from ENCODE [15] where we had eCLIP data for that RBP but no shRNA knockdown data existed (Methods 4.6). We then tested whether our SHAP values, derived from models which had never seen knockdown of the RBP, were again predictive of the sign of effect. We found weak but positive Matthew’s Correlation Coefficient (MCC) [25] values across a range of SHAP cut-offs (Table S1), suggesting that for some RBPs our models are capable of generalizing. We also looked for specific RBPs among the set with CRISPRi that performed especially well. For the 56 features (RBP binding at a position) we tested across both cell lines, 10 of them had an MCC value greater than zero. Scatterplots of the SHAP versus Δ*ψ* upon CRISPRi knockdown of the factor are shown for two features, PRPF8 at position 5 and EIF4E at position 2 (Figure 2B). For context, PRPF8 is a known component of the spliceosome with the ability to interact with 3’ splice sites (albeit it is most known for its activity at 5’ splice sites) [26–28] whereas EIF4E has only recently been linked with splicing [29–31], though its putative position-specific effects have not been investigated until now.

This analysis revealed that our modeling approach was able to broadly identify the direction of *ψ* change and existence of functional RBP binding, while being robust to multiple SHAP thresholds. Furthermore, we noticed that RBP binding at certain positions was more predictive than others.

### 2.3 Explainability (SHAP) scores identify position-specific activation and repression

We next used the SHAP values to identify general activating and repressive behavior. For each feature (RBP at a position), we calculated two summary statistics. First, we calculated the “Avg. SHAP” value across all unique binding patterns (UBPs) within our data set. To avoid confounding with the frequency of RBP binding (see Methods 4.7), we took the mean SHAP value across UBPs where that feature was bound (Methods 4.8). Second, we calculated the SHAP “importance”, which is the average of the magnitude of the local SHAP values across all UBPs (denoted as “Avg. |SHAP|”). The first statistic gives a measure of the activating/repressing effect of an RBP at a specific position, while the second provides an overall sense of how much the feature was used, regardless of sign.

In Figure 3A (left column), we show the Avg. SHAP value for each RBP at each position in both cell lines, along with a table (right column of Figure 3A) for the number of activators (Avg. SHAP > 0) and repressors (Avg. SHAP < 0). We observed many more repressors than activators at most positions across both cell lines, with the exception of positions 3 and 4, which tended to be more balanced. To delve deeper into the specific RBPs that contained the strongest position-specific activating or repressive behavior, we filtered the Avg. SHAP values across our dataset to the RBPs containing the top 10 highest and top 10 lowest values in each cell line (Figure 3B; Methods 4.9). We also provided the Avg. SHAP values across all features in both cell lines as a resource (Figure S2; Table S2).

**Figure 3.**
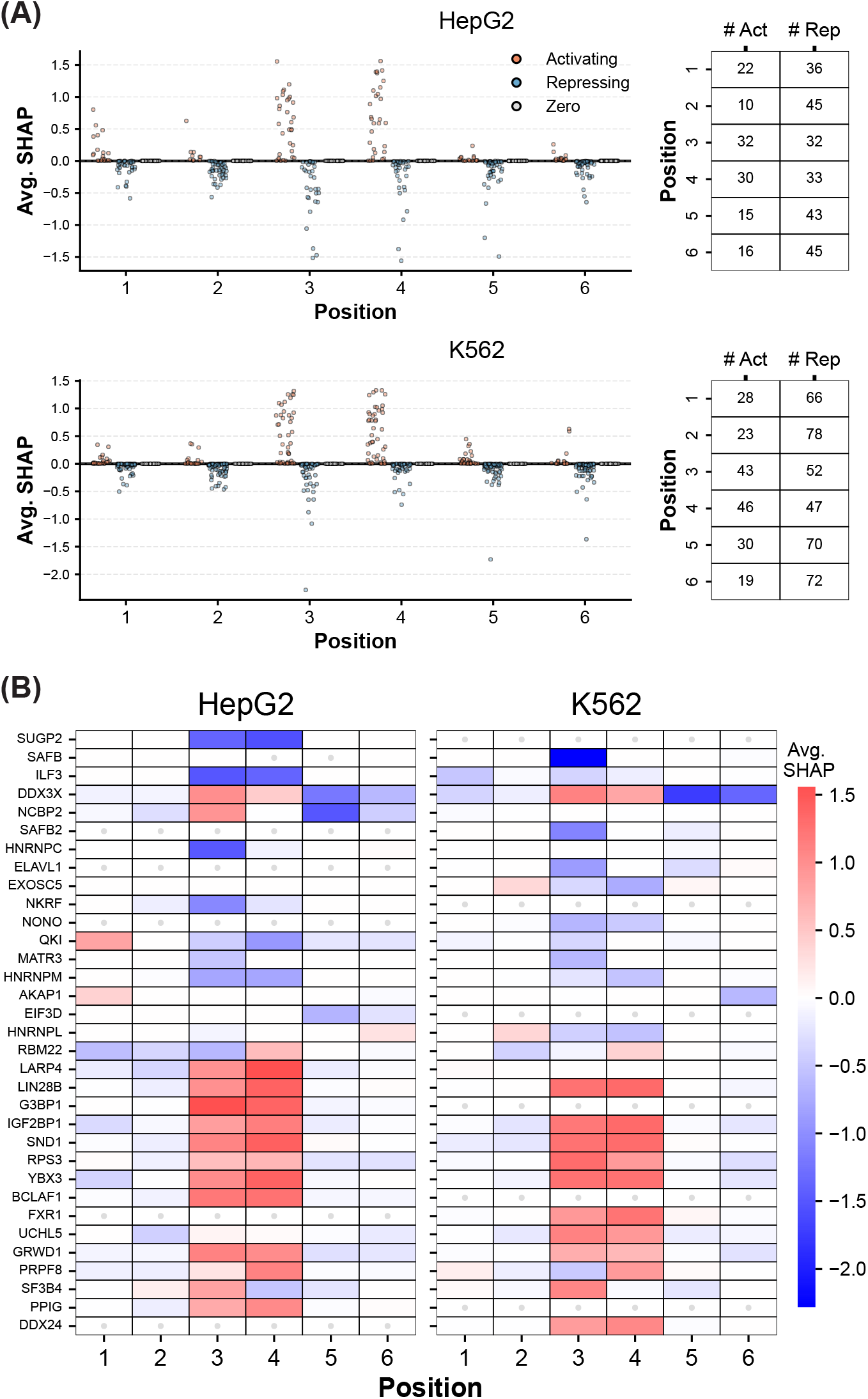
Average SHAP values indicate position-specific repressive and activating behavior of RBPs. **(A)** “Avg. SHAP” (Methods 4.8) value for all RBPs at each position (in log-odds units). Avg. SHAP values greater than 0 were deemed activating features whereas those less than 0 were determined as repressing features. **(B)** Heatmap of RBPs with the top 10 most positive or negative Avg. SHAP values (Methods 4.9). Gray circles indicate cases where eCLIP data was not available for that RBP or that RBP was never bound to that position in our dataset.

Our results both recapitulate known signatures and identify novel putative mechanisms around how RBP binding affects SE splicing at these positions. As examples of positive controls that are elucidated across both cell lines (Figure 3B; Figure S2), we highlight that PTBP1 binding at position 3 causes exon skipping [32], that HNRNPM binding around the region surrounding the middle exon caused exon skipping [33], that QKI binding to position 3 represses SE splicing (likely due to competition with SF1 over the branchpoint sequence) [34], and that the particularly strong, consistent Avg. SHAP values of DDX3X may be due to its activity specific to cancer adaptation mechanisms [35].

We also observe a number of putative novel hypotheses. ILF3 may have a role in repressive activity at position 3, in addition to previous work highlighting its repressive role at position 4/downstream of the middle exon [36, 37]. We posit that the broader biological role of MATR3 binding at position 3 may be repressive outside of the more specific contexts in which this phenomenon has been observed [38–40]. Our results also suggest a repressive role of AGGF1 binding to position 3, which has been shown in a specific context to be a repressive mechanism using a single target gene [41], may be a general regulatory rule (Figure 3B; Figure S2).

We also note that the position-specific RBP regulatory behavior discussed above was not caused by a scenario where the RBPs bound exclusively at some positions but not others. As illustrated in Figure S3, the percentage of the time an RBP binds a given position is not necessarily related to its Avg. SHAP value. This indicates that our models are picking up on patterns that go beyond simple positional preferences.

Given the noticeable trend that, across both cell lines, Avg. SHAP values at positions 3 and 4 have larger magnitudes, we hypothesized that RBP binding at the middle exon may actually be more important to understanding and explaining SE splicing than RBP binding to the flanking exons. To test this, we compared SHAP importance (Avg. |SHAP|) values from positions 3 and 4 with SHAP importance values from the other positions. We found that the middle exon positions did have high SHAP importance values, indicating that the model used them more often in making predictions (Figure S1C; Methods 4.10). Using the Avg. SHAP values, we also observed that RBP binding to middle exon positions increased *ψ* while RBP binding to all remaining positions around the flanking exons decreased *ψ* in both cell lines (Figure S1D).

In addition, we developed a model-free activator/repressor score for each feature, which we call the “Activity” score (defined in Methods 4.11). This score compares the number of statistically significant events where *ψ* increases versus decreases upon RBP knockdown and also contains a corresponding binding site (Figure 4A). An Activity score of one indicates that the RBP is always an activator at that position and a score of negative one indicates it is always a repressor. These Activity scores are visualized in Figure 4B. We observed concordance between these scores and the Avg. SHAP values for several features, such as but not limited to HNRNPC at position 3, UCHL5 across multiple positions, and PTBP1 at position 3 (Table S2). Broadly, the sign of the Activity score and Avg. SHAP were the same for 43.7% of features in HepG2 and 48.5% of features in K562 (Figure 4C). The Activity score; however, did tend to identify more features as activators, with 81 and 74 features with a positive Activity score in HepG2 and K562, respectively. This is in comparison to 37 and 27 repressive (negative Activity score) features in HepG2 and K562, suggesting that the Activity score and Avg. SHAP capture different views of RBP-driven splicing.

**Figure 4.**
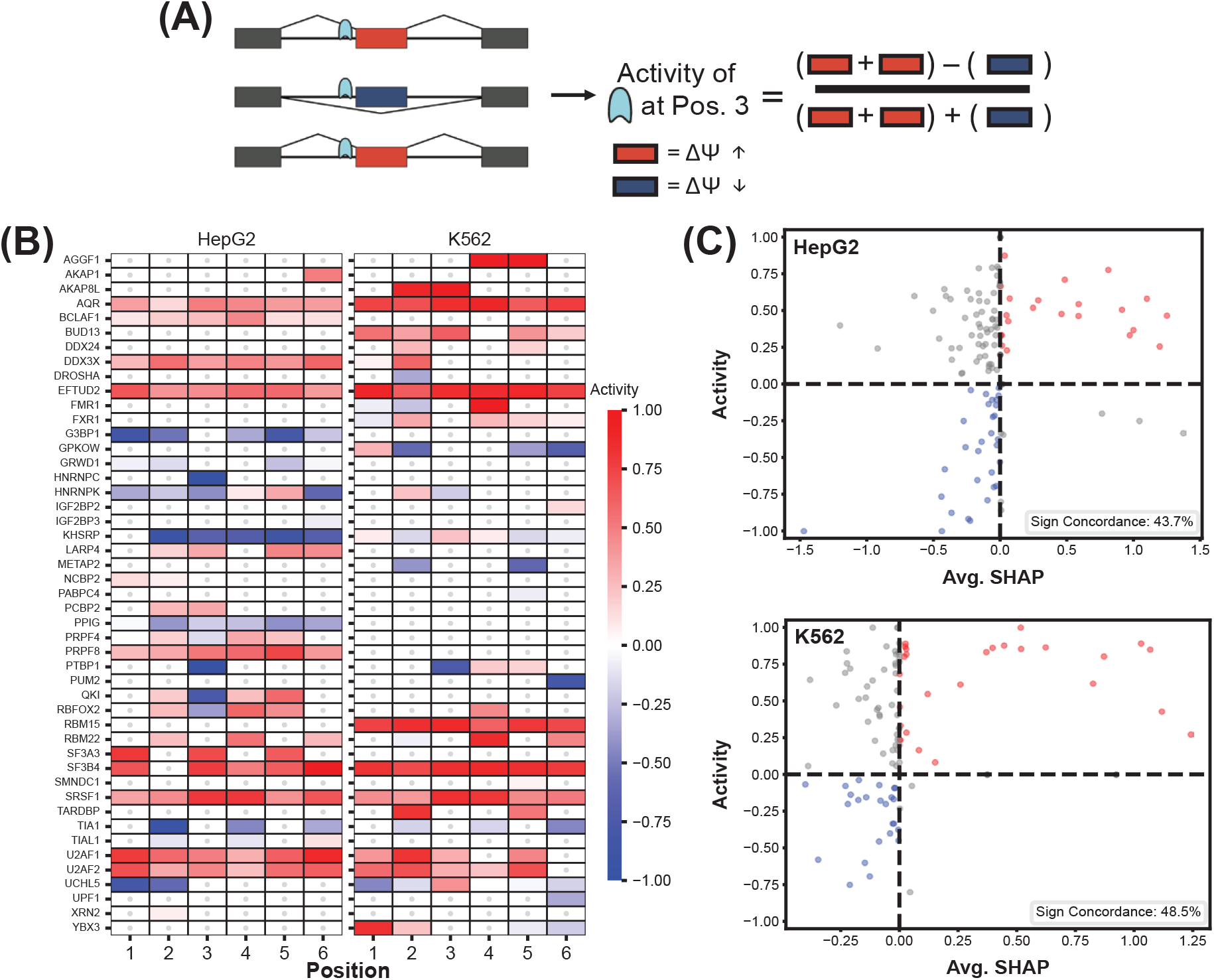
Model-free assessment of RBP-driven splicing indicates more activation than repression. **(A)** The RBP “Activity” score at a position is a normalized difference between the number of significant events where the RBP knockdown increased splicing (positive Δ*ψ*) versus decreased splicing (negative Δ*ψ*) for events with that RBP’s position-specific binding site (Methods 4.11). **(B)** Heatmap of Activity scores. Gray values indicate cases where there were not at least 10 significant events (FDR ≤ 0.1; Methods 4.11). **(C)** Scatterplot of Avg. SHAP values versus Activity scores in HepG2 (top) and K562 (bottom).

### 2.4 SHAP interaction effects reveal co-operative RBP regulation and protein-protein interactions

In addition to individual RBP effects on exon skipping, we wanted to identify co-regulatory interactions between RBP pairs at specific positions. To assess this, we took our top XGBoost model in each cell line (Methods 4.3) and generated local SHAP interaction values for all possible feature pairs (Methods 4.12). We then calculated the magnitude (importance) of a given interaction (denoted as “Avg. |Interaction SHAP|”) and the (signed) mean of these interactions (denoted “Avg. Interaction SHAP”) for all feature pairs (Methods 4.12 and Table S3).

We began by gauging the extent to which the sign (activating/repressing) was observed as being the same for feature interactions that we learned in each cell line. We show that the percentage of feature interactions with sign concordance across both cell lines is quite high and generally increases as we increased the threshold of the Avg. Interaction SHAP threshold (Figure 5A). As well, we observed that interactions between RBPs at positions 3 and 4 were the most heavily used (Figure S5A), similar to the individual effects seen in Figure S1C.

**Figure 5.**
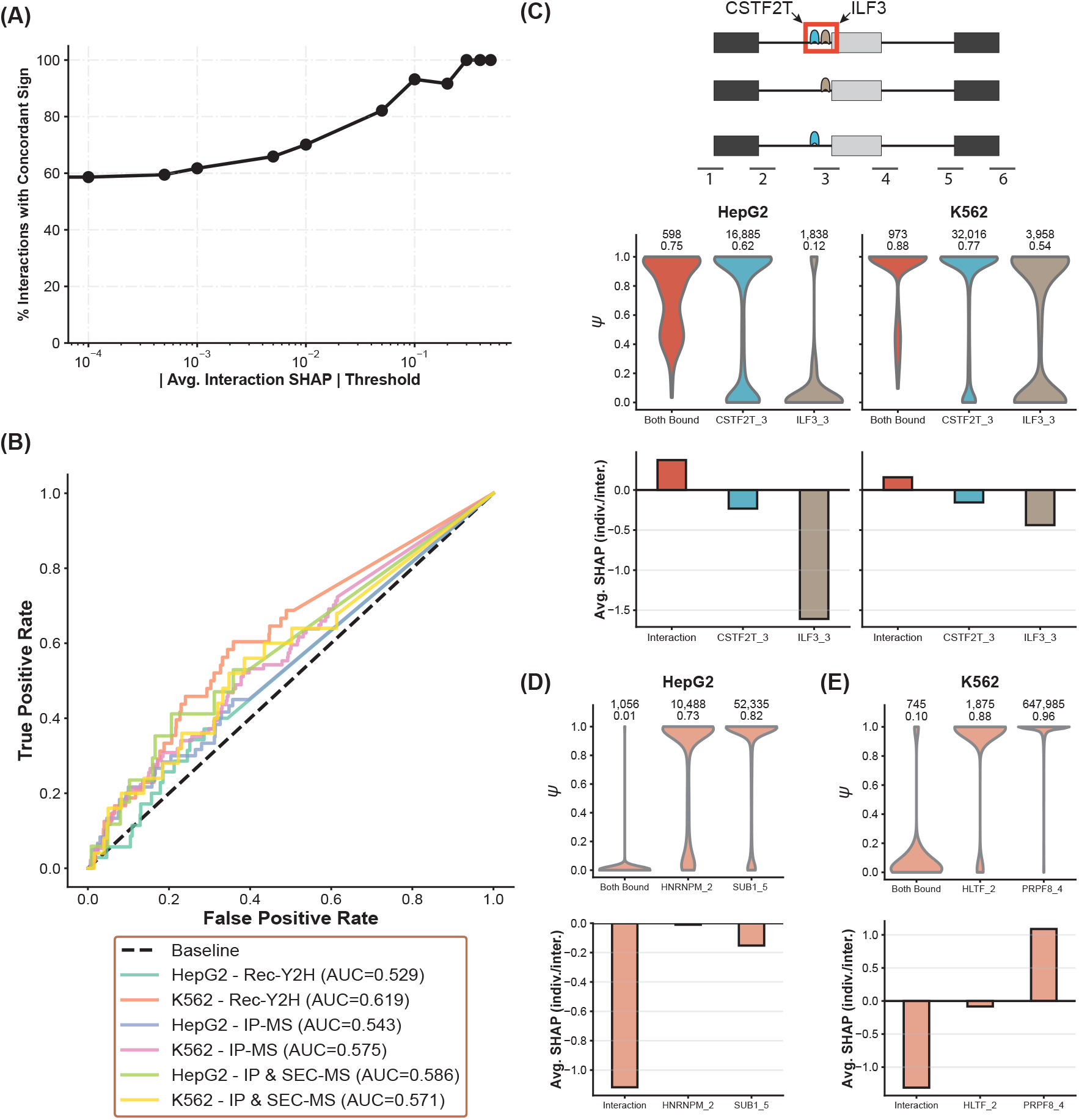
SHAP interactions capture co-regulatory roles and PPIs between RBPs. **(A)** Sign agreement between cell lines of the “Avg. Interaction SHAP” values (Methods 4.12) for each learned feature (non-zero Avg. Interaction SHAP) across different thresholds. **(B)** Receiver Operating Characteristic (ROC) and Precision Recall (PR) curves assessing Avg. |Interaction SHAP| as a predictor of PPIs between RBP pairs (details in Methods 4.14). Rec-Y2H indicates PPIs derived from ref. [42] whereas IP-MS and SEC-MS indicate PPI data from a more recent study [43] (Methods 4.13). **(C)** Example of a sign concordant interaction between CSTF2T and ILF3 at position 3. The top row displays actual *ψ* values for all SIs where CSTF2T and ILF3 were co-bound at position 3 (Both Bound) and for cases where one feature was exclusively bound (CSTF2T_3 and ILF3_3, respectively). Above each violin plot is the number (top) and the average (bottom) of the *ψ* values. The bottom row of subplots indicate the corresponding SHAP averages. See Methods 4.15 for details. **(D)** A synergistic cell-line-specific interaction between HNRNPM at position 5 with SUB1 at position 5 in HepG2 (Methods 4.15). **(E)** An antagonistic interaction between HLTF at position 2 with PRPF8 at position 4 in K562.

RBPs frequently form protein-protein interactions (PPIs) that can influence splicing [42, 43]. We hypothesized that RBP pairs which the model consistently used for prediction (as determined by the importance of our SHAP interactions) could be predictive of RBP-RBP PPIs. To test this, we used two different protein-protein interaction data sets [42, 43] to form our ground truth set of PPIs (Methods 4.13). As shown in Figure 5B, we found that the SHAP interaction importance for RBP pairs was indeed predictive of PPI status (as determined by the AUROC) across both cell lines (Methods 4.14). Given that our SHAP interaction values compare an RBP at a position with another RBP at a position (e.g., the co-binding importance of AQR at position 2 and BCLAF1 at position 5), we also checked how our results were affected by whether interactions occurred at the same position or between different positions. Generally, we found that interactions at the same position were more predictive of a PPI than pairs at different positions but that these results overall were robust to multiple different PPI resources used and were always better than random (Figure S4). Furthermore, this result replicated in both cell lines; however, we did note consistently better performance in K562 and specifically that Avg. |Interaction SHAP| values for same position PPIs were consistently stronger in K562 than in HepG2. Overall, our results shed light on how feature interactions learned by our models can decipher splicing roles for RBP-RBP PPIs.

Next, we screened our feature interactions to find places where our models learned relationships that were consistent across both cell lines and had significant actual *ψ* changes between co-binding or mutually exclusive binding events (Table S4 and Methods 4.15). We identified several candidates where RBPs co-regulate in a position-dependent manner. Whereas our previous results identified a putative novel hypothesis around ILF3 acting as a repressor at position 3 (Figure 3B and Results 2.3), we uncover a surprising activating role at this position for ILF3 when co-bound to CSTF2T at position 3 (Figure 5C). CSTF2T has well-documented roles in 3’ end processing, suggesting novel potential biological interactions between splicing and 3’ end processing [44, 45]. We also note other interesting vignettes, found in both cell lines, such as the case of co-binding by DDX3X at position 2 with PTBP1 at position 3 (each discussed in Results 2.3) reversing the known repressive role [32] of PTBP1 at that position (Figure S5B) and a case whereby the co-binding of QKI at position 3 (as discussed in Results 2.3) with U2AF2 at position 3 reverses the broad activator role [46] of U2AF2 (Figure S5C).

Finally, we also focused on feature interactions where the interaction effect was stronger than any of the individual effects. Specifically, we looked for cases where the actual *ψ* distributions were significantly different between the co-binding examples and the exclusive individual binding examples (Table S5; Methods 4.15). We did not enforce any requirements on concordance between cell lines as we used this as an opportunity to highlight certain novel putative cell-line-specific interaction effects. As a result, we found a putative interaction with a synergistic effect between HNRNPM at position 2 and SUB1 at position 5, leading to prominent exon skipping (Figure 5D). HNRNPM is a documented splicing repressor [47, 48] whereas the role of SUB1 is less known [49, 50], albeit initial evidence indicates it may have an effect on RNA expression of its targets [50]. Specific to the K562 cell line, the binding of HLTF at position 2 with PRPF8 at position 4 reverses the activating role of PRPF8 at this position and causes a decrease in *ψ* (Figure 5E). While PRPF8 is well-known in affecting splicing by being part of the spliceosome [27, 51–53], HLTF is known for its various roles including but not only limited to maintaining genome integrity [54, 55] and activating transcription of oncogenes [56]. Yet, no role has currently been found for HLTF in splicing, potentially suggesting a novel function for this protein within blood cancer contexts.

## 3 Discussion

The results of this study aim to address how functional patterns of RBP binding lead to exon skipping. By jointly modeling the binding of over 168 RBPs, we sought to disentangle both the individual and collective effects of position-specific RBP binding on exon skipping. To do this, we first integrated RBP eCLIP data with RBP knockdown followed by RNA-seq data from the HepG2 and K562 cell lines. In training models to predict *ψ* for an exon skipping event based on the position-specific RBP binding of that event, we found that XGBoost, a more complex machine learning method, outperformed a simpler elastic net model. This success indicates that more sophisticated machine learning architectures that integrate additional binding [57, 58] and splicing information [11, 12, 14, 59–62] may further improve our understanding of the relationship between RBP binding and splicing.

To better understand what, if any, regulatory logic the XGBoost models learned, we applied SHAP, a machine learning explainability technique, to determine how much the model used each binding site in a given prediction. Using these values we were able to build up a transcriptome-wide picture of the activating and repressive nature of each RBP around different exon boundaries (positions) and found that the model preferentially used the splice sites surrounding the middle exon. While we provided the models with the position of each binding site, they were unaware of relationships between positions, indicating that this positional preference for the middle exon was discovered by the model *de novo*. In addition, interactions between RBPs near the 5’ and 3’ ends of the middle exons were used more than others.

To interrogate the effect of removing an RBP binding site on *ψ*, we compared the local SHAP value for a feature (RBP bound at a position) with the change in *ψ* upon knockdown of that RBP. Using exon skipping events unseen by the model, we found that the sign of the local SHAP for an RBP binding site was predictive of the increase/decrease in *ψ* upon knockdown of that RBP. Furthermore, for a set of RBP knockdowns not in our original data set, we found broad sign concordance between local SHAP sign and change in *ψ* upon RBP knockdown. This result, in particular, suggests that our models and the logic they have learned are not dataset-dependent.

We found that our SHAP-estimated activation/repression showed reasonable agreement when compared to differential splicing events where the knockdown RBP was bound (the “Activity” score). However, giving the limited number of differential splicing events with RBP binding sites, our average SHAP estimates were able to provide a broader picture of activation and repression. We also found that the average SHAP metrics revealed RBP-specific position preferences for “functional” binding (i.e., those binding sites likely to impact *ψ*). For example, we recapitulated known signatures for the RBPs PTBP1, HNRNPM, QKI, DDX3X, and more, while identifying putative novel position-resolved hypotheses for ILF3, MATR3, and AGGF1.

Alternative splicing is known to require the coordination of many RBPs, which leads to the question of which RBPs work together to effect splicing in a position-specific manner. To identify these putatively co-regulatory effects, we calculated SHAP feature interactions for pairs of RBPs across positions. We found that most of these average interaction values were highly concordant between cell lines, suggesting robust estimates. Furthermore, the average SHAP interaction was predictive of protein-protein interactions between RBPs. This result would support the use of PPI as an embedding to more sophisticated models of splicing prediction. In particular, this may be of use for predicting the effect of unseen perturbations, as models trained to predict the gene expression response to a perturbation see improved performance when adding PPI embeddings [63]. We also found a number of position-dependent effects between RBPs where the presence of both RBPs switched the impact of binding from increasing *ψ* to decreasing or vice versa. Further experiments would be needed to validate these “sign-switching” effects; however, our interaction estimates provide promising (and testable) hypotheses.

In conclusion, we find that machine-learning models can successfully predict *ψ* from experimentally-derived RBP binding patterns. We also find that the logic learned by these models is biologically interpretable and can be used to interrogate the effects of RBP binding on exon skipping. Our work provides an example of how machine learning models designed to provide functionally testable hypotheses can advance biological knowledge. We anticipate that modeling in this way will help move machine learning in genomics from prediction to scientific understanding.

### Study Limitations

While our RBP modeling effort is one of the largest to date in terms of number of RBPs considered, our ENCODE-derived eCLIP data contained binding sites for 168 RBPs out of the thousands of known RBPs [64, 65]. In addition, our model considers only eCLIP binding, which can be noisy and suffers from exonic bias [15, 19]. Future modeling efforts may benefit from integrating both eCLIP and RBP motif scans. Another important consideration is our choice of including binding events within a given window around exon boundaries. For this work, we found that our models performance decreased as we extended the window size. Thus, we faced a trade-off between considering more binding sites in our SHAP analyses versus model performance.

## 4 Methods

### 4.1 Integrating ENCODE shRNA RNA-seq and eCLIP data

The ENCODE portal [66] was used to download all eCLIP [17] peak BED [67] files and shRNA RNA-seq (shRNA) FASTQ data for the HepG2 and K562 cell lines [15]. The eCLIP dataset was chosen by selecting data from the HepG2 and K562 cell lines and taking all GRCh38 genome assembly released narrowPeak BED files that came from the IDR process [68]. For eCLIP RBPs that had 2 experiments released concurrently, we took the newest experiment. In total, we obtained 105 RBPs with eCLIP and 85 RBPs with matching shRNA knockdown RNA-seq in HepG2. In K562, we downloaded eCLIP data for 139 RBPs, 101 of which had shRNA knockdown RNA-seq. We made sure to only use shRNA experiments that selected for polyadenylated mRNA in this work.

#### 4.1.1 RNA-seq pipeline and rMATS splicing analysis

To investigate alternative splicing differences between the RBP knockdown and control samples, we employed a comprehensive short read RNA-seq pipeline. Prior to alignment, we prepared a composite reference genome and transcriptome annotation. We obtained the GRCh38 no-alt analysis set primary assembly (GCA_000001405.15) and appended two ERCC spike-in FASTA files (ENCFF001RTP and ENCFF335FFV) to the primary genome sequence. For gene annotation, we used the GENCODE V29 primary assembly GTF, to which we concatenated the GENCODE V29 tRNA annotations and custom GTF entries generated from the spike-in FASTA headers. This produced a single combined FASTA and a single combined GTF that were used for all subsequent index generation, alignment, and splicing quantification steps. A STAR genome index was then generated from these combined files.

For each shRNA knockdown experiment, paired-end FASTQ files were retrieved programmatically from the ENCODE portal using the ENCODE REST API. Given an experiment accession identifier, the pipeline queried the ENCODE API to resolve the accession into individual FASTQ file accessions, determined which files correspond to knockdown versus control samples by inspecting the “possible_controls” field of the experiment metadata, and correctly paired forward and reverse reads using the “paired_with” annotation. FASTQ files were downloaded for all matched RBP knockdown and control samples in the HepG2 and K562 cell lines.

Paired-end reads were aligned to the composite reference genome using STAR (version 2.7.11a) [69] in two-pass mode (“twopassMode Basic”). Key alignment parameters included local alignment (“alignEndsType Local”), a maximum of 4 mismatches (“outFilterMismatchNmax 4”), splice junction overhang minimums of 5 base pairs (bp) and 1 bp (“alignSJoverhangMin 5”, “alignSJDBoverhangMin 1”), an intron size range of 20 to 1,000,000 bp, and output filtering by splice junction (“outFilterType BySJout”). The modified and composite V29 GTF was supplied via “sjdbGTFfile” for splice junction annotation. Output was written as coordinate-sorted BAM files. The same STAR index and parameter configuration were used for all samples across both cell lines.

BAM files from the knockdown and corresponding control samples for each RBP were then provided to rMATS-turbo (version 4.2.0) [23] for differential splicing analysis. rMATS was invoked in paired-end mode (“-t paired”) with a strand-specific library type (“libType fr-firststrand”), a nominal read length of 100 bp, and with the “variable-read-length”/”--allow-clipping” flags enabled to accommodate heterogeneity in read lengths across the RNA-seq samples. The composite GTF file (provided during STAR alignment) was again supplied for transcript annotation. rMATS-turbo quantifies alternative splicing in terms of Percent Spliced In (*ψ*) [70], defined as the ratio of the Inclusion Junction Count (IJC) to the sum of the IJC and Skipped Junction Count (SJC). Junctions for which both IJC and SJC were zero are excluded. We defined the change in Inclusion Level from knockdown (sample 1 in rMATS-turbo) to control (sample 2 in rMATS-turbo) as the quantification of differential alternative splicing. The significance of the difference in Inclusion Levels was calculated by rMATS-turbo using a maximum likelihood estimate to compute the significance expressed as a corrected p-value corresponding to each alternative splicing event. rMATS-turbo reports dysregulation across five modes of alternative splicing: skipped exon (SE), mutually exclusive exons (MXE), alternative 3^′^ splice site (A3SS), alternative 5^′^ splice site (A5SS), and retained introns (RI). For this study, we used the SE results.

Quality control was performed at both the alignment and splicing quantification stages. STAR alignment summary files (“Log.final.out”) were parsed for each sample to extract metrics including the number of input reads, the percentage of uniquely mapped reads, the total number of detected splice junctions, and the percentage of multi-mapped and unmapped reads. Samples with fewer than 70% uniquely mapped reads were flagged for review. rMATS summary files were similarly collected and inspected across all experiments.

### 4.2 Integration of eCLIP data with rMATS output

In order to model an exon skipping event’s *ψ* as a function of RBP binding, we integrated the eCLIP data described above with the rMATS output file for SE splicing as follows. First, for every exon skipping event observed in any of the rMATS knockdown files processed above, we mapped all eCLIP peaks to the closest exon boundary and retained those peaks that were within 100 bp of the 5’/3’ ends of each boundary, which we labeled positions 1-6 (Figure 1A). Peaks equidistant from exon boundaries were randomly assigned. We only kept peaks and exon boundaries from the human autosomal chromosomes. Thus, for each exon skipping event observed in our data, we had a list of RBPs with binding sites around each position. We then created a table where each column is a feature representing a specific RBP at a specific position (e.g., QKI at position 5) and each row represented the coordinates of an exon skipping event for a given condition. A “1” indicated “binding” of that RBP at that position, and “0” indicated “no binding”.

Second, we merged the *ψ* value for each exon skipping event in a given condition (control/knockdown) with the corresponding binding table above. Each RBP knockdown experiment from ENCODE contained two knockdown RNA-seq samples and two control samples, resulting in four *ψ* measurements. For exon skipping events coming from an RBP knockdown sample, we applied an “in-silico knockdown”, where we switched a “1” in our table to a “0” in all positions where that knockdown RBP was bound. We then repeated this for all RBP knockdowns in our data (85 in HepG2 and 101 in K562). We excluded rMATS events with fewer than 40 total reads across the inclusion and skipping junctions in rMATS. After integrating all knockdown and control RNA-seq samples with the binding data, we have a table where each row contains the coordinates of the three exons involved in an exon skipping event, the position-specific binding around that event, and the corresponding *ψ* value. We define this as a “splicing instance” (SI). Our final datasets contained approximately 4.57 million SIs in HepG2 and 6.29 million in K562. Since the ENCORE experimental design involved shared controls (i.e., multiple RBP knockdown experiments compared against the same control RNA-seq experiments), we de-duplicate the data so that the same SE event from the same control RNA-seq sample does not show up more than once.

### 4.3 Model training and evaluation

To train and evaluate the elastic net and XGBoost models, we employed the following strategy. For each cell line, we took all SIs (i.e., rows in our dataset) and grouped them by the gene they came from. Next, we randomly split the data into 5 folds with an approximately equal number of genes in each fold. A single fold (20% of genes) was taken as the final holdout set for that cell line whereas the remaining 4 folds (80% of the genes) were used for training and validation. We then split the 4 training folds into 5 inner folds (by gene) and evaluated the predictions across all 5 splits, where each fold had a chance to be the test data (holdout *R*^2^). This inner fold splitting was repeated with 7 random seeds, where the holdout *R*^2^ values were averaged across the seed splits to evaluate performance. All models and model comparisons were performed separately for each cell line.

#### 4.3.1 Elastic net and XGBoost modeling

We applied elastic net linear regression models to our datasets as a baseline. The binarized binding (“0”/”1”) of each RBP at a specific position was used as a term (i.e., feature) in the regression, with the corresponding *ψ* value as the response variable. To ensure comparability between the elastic net and XGBoost models, we made sure to utilize the same training/test splits for both. We tuned the elastic net model in each cell line separately and used 7 training seeds per hyperparameter configuration to retrieve our averaged holdout *R*^2^ score, with the best elastic net model hyperparameters chosen based on the maximum average.

For our XGBoost models, we again used the binarized positional binding of the RBPs as the inputs to the XGBoost framework [24] (Python package version 2.1.1) and trained models to predict the corresponding *ψ* value. We took a superset of the hyperparameters that are generally tuned when applying XGBoost to real-world data [71–76]: namely, “max_depth”, “learning_rate”, “min_child_weight”, “gamma”, “subsample”, “colsample_bytree”, “reg_alpha”, and “reg_lambda”. We used a 100 round threshold for early stopping (10% of training data was used as validation set) and set our “n_estimators” value to an arbitrarily large number (100,000) along with using the “reg:logistic” objective. For the remaining hyperparameters, as a full grid search was computationally prohibitive, we took a sequential tuning approach that aimed to choose the best hyperparameters (one set of them at a time) by maximizing the averaged holdout *R*^2^ values. For details, please refer to the corresponding GitHub repository. For our final holdout (“test”) dataset performance evaluation as well as for XGBoost-derived SHAP calculations, we took the top 5 XGBoost and top 5 elastic net hyperparameter configurations based on each configuration’s averaged holdout *R*^2^ score in a cell-line-specific fashion.

### 4.4 Calculating local SHAP values

We retrieved SHapley Additive exPlanation (SHAP) [22] values for every SI in our dataset (i.e., all training, validation, and test rows) using the “TreeExplainer” [77] function in the Python “shap” package (version 0.49.1), with the local SHAP values being in log-odds units. The term “local” implies a SHAP value for a feature within one SI. For deeper intuition, please see Methods 5.1. To account for variability in model runs, we averaged the local SHAP values across our top 5 performing models in each cell line, respectively. For all analyses, we consider local SHAP values only for places where that feature was bound in that SI.

### 4.5 Concordance between Δ*ψ* and local SHAP

We used the rMATS differential splicing results for an RBP knockdown to retrieve the change in splicing (Δ*ψ*) for rMATS events (FDR ≤ 0.1) from the test data of a given cell line. We subset these rMATS events to cases where the RBP was bound at any position in the corresponding control sample. We then took the local SHAP value for this RBP and position feature in the event’s corresponding control SI. Next, we compared the sign of the Δ*ψ* against the sign of the local SHAP using Matthew’s correlation coefficient (MCC) [25]. The Δ*ψ* values here were multiplied by -1 to be in the order of control and then knockdown meaning that positive Δ*ψ* indicated “activator” whereas negative indicated “repressor”.

### 4.6 CRISPRi RBP knockdown external validation data analysis

In order to assess concordance between Δ*ψ* and local SHAP values in an external dataset, we integrated CRISPRi knockdown RNA-seq from ENCORE [15] for RBPs that we did not have shRNA knockdowns for in a given cell line, but did have eCLIP.

We processed the CRISPRi knockdowns very similarly to our shRNA RNA-seq knockdown data (Methods 4.1.1), including using the same composite GTF file as input for alignment and splicing quantification, except we utilized rMATS-turbo version 4.3.0 and STAR version 2.7.11b. We created SIs for the SE events coming from these knockdowns by matching the six exon boundary genomic locations of these events with the same locations of SE events in our main shRNA-derived SIs, in a cell-line-specific fashion. Because the event locations are identical, the binding pattern is identical and hence, the local SHAP values for a given SI will be equivalent. Knowing this, we subsetted to events where a given RBP at a position was in-silico knocked down (since it came from an RBP knockdown sample and those genomic coordinates originally contained the feature’s binding). We only considered events where at least one sample in each condition had more than 10 reads. Next, we took the Δ*ψ* and rMATS FDR value from the CRISPRi RBP knockdown rMATS file to compare against the local SHAP value for that feature. Similar to Methods 4.5, we reversed the sign of the Δ*ψ* values. After, we calculated the Matthew’s correlation coefficient between Δ*ψ* and local SHAP for each feature only considering SE events with an rMATS FDR value ≤ 0.1. Independently, we also concatenated all aforementioned per-feature examples to calculate MCC values across the CRISPRi dataset, while using multiple minimum thresholds for the SHAP value per example.

### 4.7 Defining unique binding patterns (UBPs)

Different exon skipping events across the transcriptome can have the same binding pattern. Because our model only uses an RBP binding pattern as its input, it will predict the same *ψ* and yield the same SHAP values. To avoid inflating our SHAP statistics, SHAP metrics were summed over the set of unique binding patterns (UBPs) in a cell-line specific fashion. The number of UBPs per cell line was approximately 115,000 and 148,000 for HepG2 and K562, respectively. Importantly, when comparing SHAP-derived quantities to the observed *ψ* or Δ*ψ*, we instead used the local SHAP for that corresponding SI.

### 4.8 Calculating global feature behavior metrics

We sought to calculate two statistics based on local SHAP values. First, what is the average magnitude of effect (“importance”) of an RBP at a given position across all its binding events? To do this, we took the average of the absolute value of the local SHAP values for that feature (RBP at a position) across all UBPs (Methods 4.7) where that feature was bound, which we denote as “Avg. |SHAP|”. Second, we wanted an estimate of both the magnitude and direction, on average, of a feature to indicate whether it was an activator or a repressor. For this we simply took the (signed) mean of the local SHAP values for a specific feature, once again over the UBPs where that feature was bound and denote this value as “Avg. SHAP”.

### 4.9 Vignette heatmap of strongest activator and repressor behavior

We subsetted to the RBPs with the top 10 highest and lowest “Avg. SHAP” values in each cell line. For RBPs that appeared in the “highest” category at one position, but “lowest” at another position, we assigned those RBPs to the “lowest” section of the plot. Light gray dots denote places where either that RBP did not have eCLIP data or where no examples existed of that RBP binding to that position.

### 4.10 Comparing Avg. |SHAP| of positions 3 and 4 versus all other positions

For each cell line, we took the “Avg. |SHAP|” values per feature and grouped them based on whether they were from positions 3 or 4 versus if they were from any of the other positions (i.e., 1, 2, 5, 6). We ran a one-sided Mann-Whitney U test to see if the distribution of Avg. |SHAP| values for positions 3 and 4 were stochastically larger (“greater”) than for positions 1, 2, 5, and 6.

### 4.11 “Activity” score

We assessed the splice activating or repressing activity of an RBP in a position-specific manner based on the rMATS differential splicing (Δ*ψ*) estimates for events where the knockdown RBP had a binding site.

Specifically, we took a ratio comparing the number of significant differential SE events where *ψ* was decreased upon knockdown versus where *ψ* was increased upon knockdown, defined as the “Activity” score (see equation below).

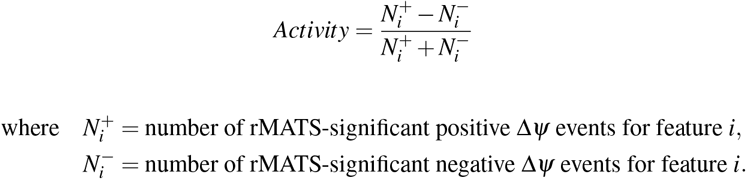

In this analysis, we reversed the sign of the Δ*ψ* value so that positive values corresponded to activation and negative values corresponded to repression (as done in Methods 4.5). We restricted our analyses to features where there were at minimum 10 rMATS-significant splicing events (rMATS FDR value ≤ 0.1) where that feature was bound.

### 4.12 Feature-feature interaction SHAP analysis

Similar to an interaction term between two variables in a linear model, we can calculate a SHAP equivalent for XGBoost. Specifically, we calculate local interaction SHAP values for all pairs of features. As discussed in the individual effect local SHAP value generation (Methods 4.4), all the local SHAP values for a given row add up together to the difference between the model predicted value (in log-odds units) and the “baseline” value. Hence, for the interaction effect local SHAP, the SHAP values for each individual feature plus the feature-feature interactions add up to the difference between the model predicted value and the “baseline” (average model prediction). Due to the large number of interactions, we restricted our analysis to the best performing XGBoost model within each cell line. Interaction values were calculated using the Python “shap” package’s “shap_interaction_values” function (version 0.49.1) as part of the “TreeExplainer” class with the options “feature_perturbation = tree_path_dependent” and “model_output = raw” applied.

Similar to the individual feature behavior calculations (Methods 4.8), we calculated the “importance” of a feature interaction (denoted as “Avg. |Interaction SHAP|”) as well as both the magnitude and direction of these interactions (denoted as “Avg. Interaction SHAP”). We calculated these two metrics using only UBPs (Methods 4.7) where both features involved in the interaction were co-bound.

### 4.13 RBP-RBP PPI data resources

We used two protein-protein interaction (PPI) data sets to create four variants of RBP-RBP PPI designations that we tested against our SHAP interactions. Our first data source was a recombination Yeast two-hybrid [78] RBP screening resource (Rec-Y2H) [42], where we used their Supplementary Table S1 and took all pairs of RBPs that they screened as “baits” or “preys”. Using their Supplementary Table S2, we subsetted their screen results table to where the “SumIS” value was greater than or equal to “7.1”, as recommended by the authors, to obtain “true” PPIs. RBP pairs that were tested but had no interaction were considered “false” in our downstream analyses, while pairs that were not tested were considered “null” values.

The second data source employed size-exclusion chromatography mass-spectrometry (SEC-MS) and immunoprecipitation-mass spectrometry (IP-MS) to understand PPIs between RBPs [43]. We used their Supplemental Table S1 to retrieve the “baits” used for IP-MS, which would tell us which RBPs were even tested for PPIs. Next, we took their Supplemental Table S2 containing the interactions they found and used the “Interaction_support” column to determine whether the interaction was found via IP-MS or whether the interaction was found in both IP-MS and SEC-MS. The “true” PPI, “false” PPI, and “null” values mentioned in the previous paragraph were similarly defined for this data source.

### 4.14 Predicting RBP-RBP PPIs from “Avg. |Indteraction SHAP|”

We tested whether the importance of SHAP interactions (“Avg. |Interaction SHAP|”; Methods 4.12) was predictive of PPIs between RBPs, using the “scikit-learn” [79] functions “roc_curve”, “roc_auc_score”, “precision_recall_curve”, and “auc” functions (version 1.8.0) to get the necessary values that went into the final visualization. We started with Avg. |Interaction SHAP| values between RBP pairs with occurrences of cobinding and subset our remaining model interaction effects to only include those where the two RBPs involved were explicitly tested (the PPI status was either “true” or “false”; details in Methods 4.13).

Given that the PPI data does not include positional RBP-RNA binding information, we next filtered interaction values to those involving RBP pairs that were bound at the same position (“Same-Position PPI”), different position (“Different-Position PPI”) or any position (“All-Positions PPI”). For each of these three position types, we summed all remaining Avg. |Interaction SHAP| values for a given RBP pair, which were then ranked from highest to lowest to render our ROC and PR curve visualizations.

### 4.15 Identifying feature interactions with significant *ψ* changes

We took all non-zero Avg. Interaction SHAP values in both cell lines and were sign concordant. Next, we subset these to where the Avg. SHAP values of the two individual features making up the interaction were also each cell line sign concordant. Importantly, the Avg. SHAP mentioned in the previous sentence is calculated from the individual feature local SHAP values generated during the creation of the interaction local SHAP values.

With this set of interaction features (and their corresponding individual features), we compared the true *ψ* distributions for where the two features were co-bound to the corresponding *ψ* distributions where they were mutually exclusively bound (in a cell-line-specific manner) and ran two one-sided Mann-Whitney U tests comparing the co-bound *ψ* distribution against each of the other two. The “side” of the test was determined by the sign of the Avg. Interaction SHAP. We only kept cases where all test p-values were less than 0.05. For our examples showcasing cell-line-specific interaction effects, this same process was employed except we did not enforce any cell line concordance requirements.

## Acknowledgements

The authors would like to thank the members of the Platig lab for helpful discussion. JP acknowledges funding from NHLBI (K25HL140186, R01HL176576). PJC received funding from NHLBI (R01HL124233 and R01HL166992). JP, PJC, JD, and AP received funding from R01HL171213. AP received funding from the Harold Alfond Foundation.

## Author Contributions Statement

YR, AP, RA, SK, AI, and JV processed and analyzed the data. YR, AP, and RA analyzed model results. AP, BJ, PJC, and JP developed the machine learning framework. YR and JP wrote the manuscript. All authors reviewed the manuscript. JD, PJC, and JP provided support for the work.

## Competing Interests

PJC received consulting fees from Genentech and Verona Pharma and grant funds from Sanofi and Bayer. AP is on the Scientific Advisory Board of Duet Biosystems and receives consulting fees from Feromics Inc. and PriveBio Inc. The other authors declare no competing interests.

## 5. Supplementary Information

### 5.1 Calculating SHAP values

To understand the regulatory impacts of RBP binding at specific exon boundary positions on SE splicing, we retrieved per-data-row (local) SHAP [22] values for every single SI in our dataset (training, validation, and test). This was done through the Python “shap” package (version 0.49.1) “TreeExplainer” [77] class with the “shap_values” function called using the options “feature_perturbation = tree_path_dependent” and “model_output = raw”. Importantly, this means that all our local SHAP values are in units of log-odds.

Put briefly, the SHAP method aims to answer, for each row in a dataset, why a given model predicted the value that it did as a function of how much each feature’s value for that row contributed to the difference between the prediction for that row and a “baseline” value that is the best approximation of the data if one had no feature information.

For example, if the predicted *ψ* value for a SI was 0.9 and the baseline value was 0.75 for the training dataset used to create the model, SHAP would provide values for every single feature in our dataset to explain how the binding or lack thereof increased or decreased the prediction such that the sum of all of these local SHAP values would equal the difference of 0.9 and 0.75 (i.e., 0.15).

For all analyses, we only considered local SHAP values for features where that feature was bound in that SI. For example, if a given SI had RBFOX2 binding to position 4 but it did not bind to position 5, the local SHAP value for RBFOX2 at position 5 was not needed or used within our work.

## 6 Supplementary Figures

**Figure S1.**
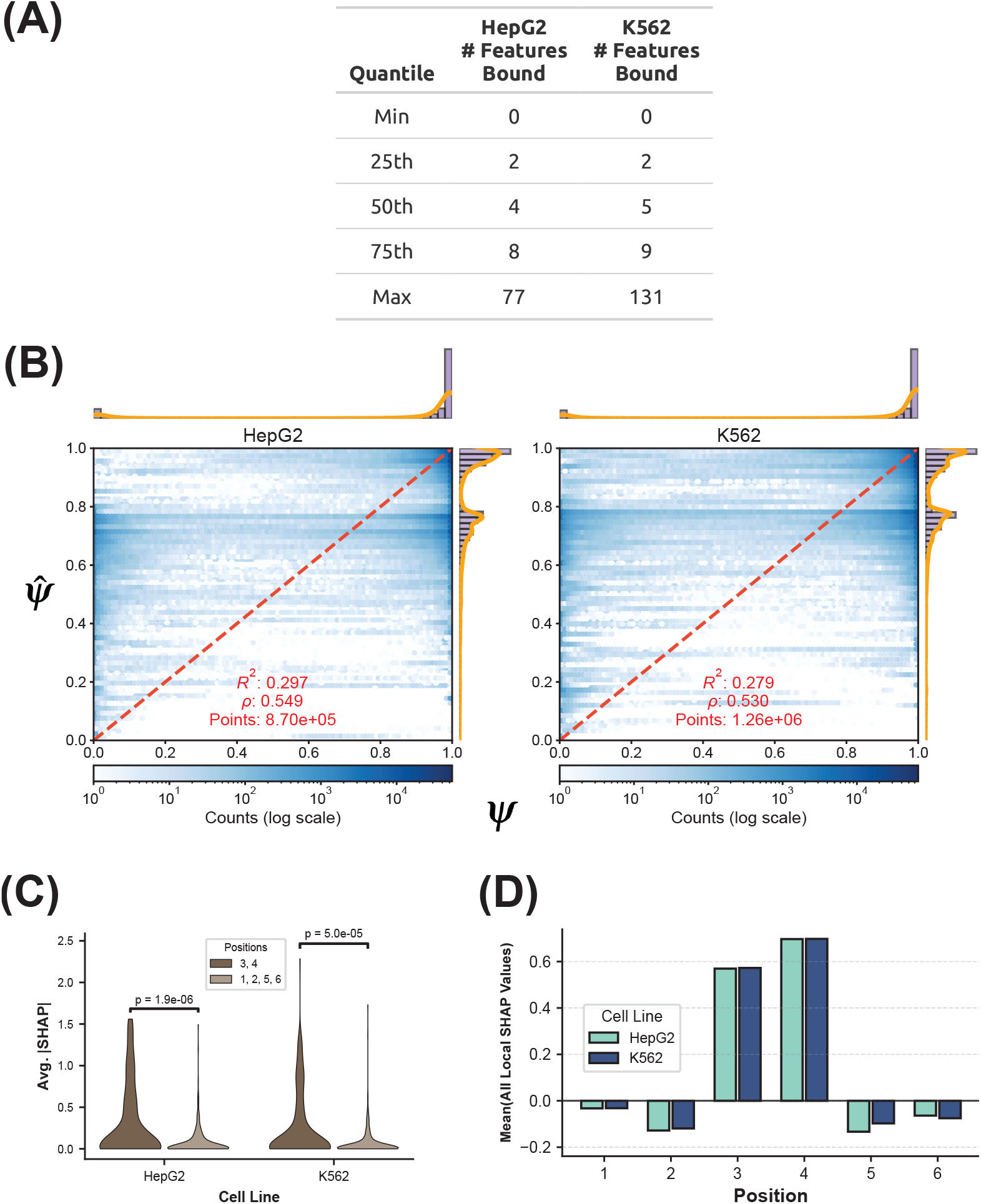
**(A)** Quantiles for the number of features bound per Splicing Instance (SI) in our dataset (Methods 4.2). “Feature” represents an RBP and position (e.g., RBFOX2 at position 4) and SIs with “0” bound features are due to cases where we did an in-silico knockdown and no other binding remained (Methods 4.2). The median number of bound features is four in HepG2 and five in K562. **(B)** Scatterplot of the actual (*ψ*; x-axis) with the predicted (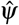 ; y-axis) values on the final holdout (“test”) dataset for the top-performing XGBoost model per cell line (Methods 4.3). The marginal distribution of *ψ* is shown along the top of the scatterplot and the marginal for 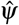 is shown along the right side. **(C)** Violin plots of “Avg. |SHAP|” values (Methods 4.8) for all features, stratified by position. A one-sided Mann-Whitney U test was run comparing whether the distribution of Avg. |SHAP| values from positions 3 and 4 was greater than the distribution from the other positions (Methods 4.10). **(D)** Mean of local SHAP values (Methods 4.4) across positions using the “Unique Binding Patterns” (UBPs, Methods 4.7).

**Figure S2.**
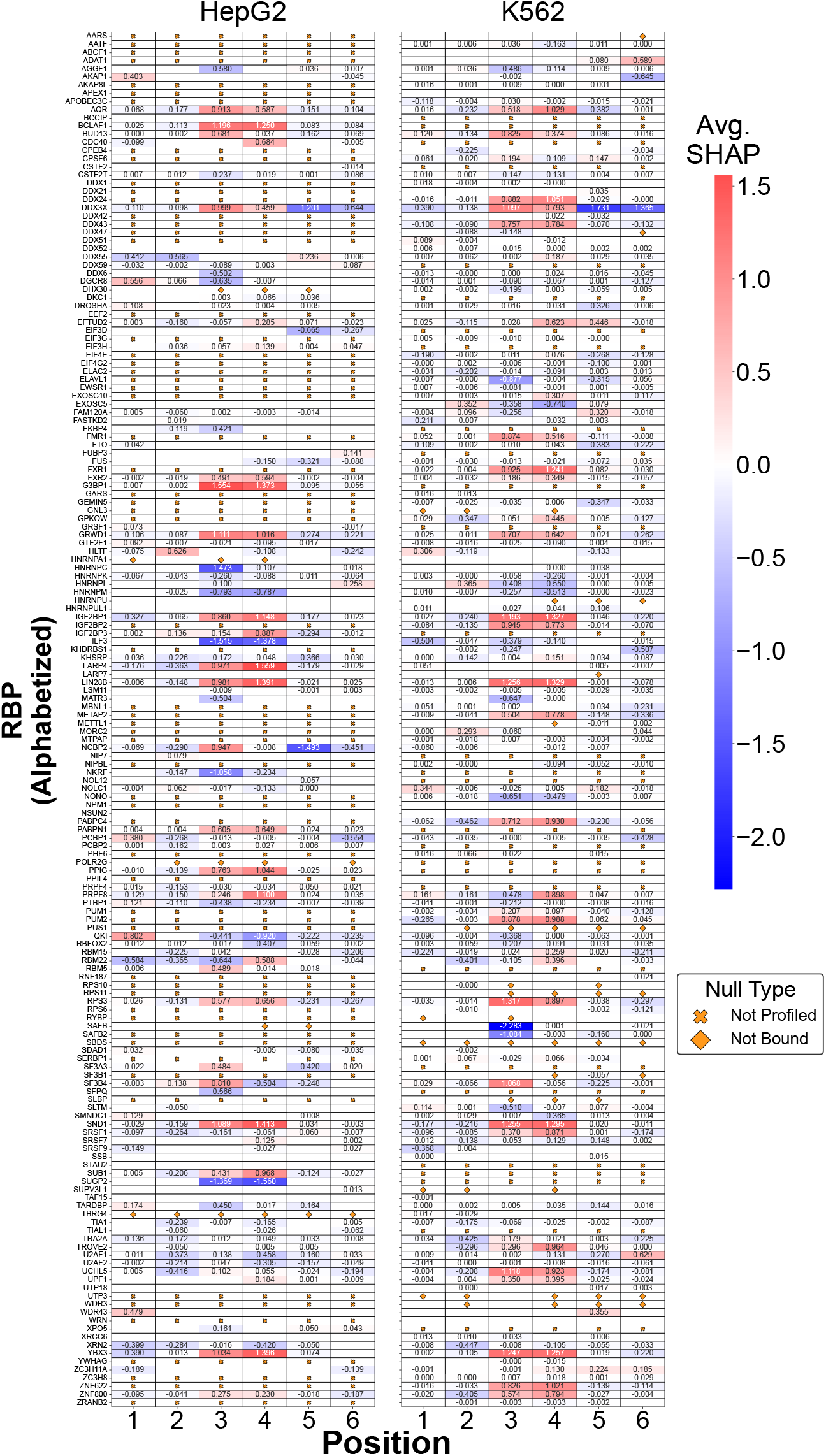
All “Avg. SHAP” (Methods 4.8) values across both cell lines, shown in log-odds units. The “×” marker represents RBPs not profiled with eCLIP in a given cell line whereas the “◊” indicates features with no binding examples. Cells without any text represented cases where binding existed but the feature had no effect (Avg. SHAP value = 0).

**Figure S3.**
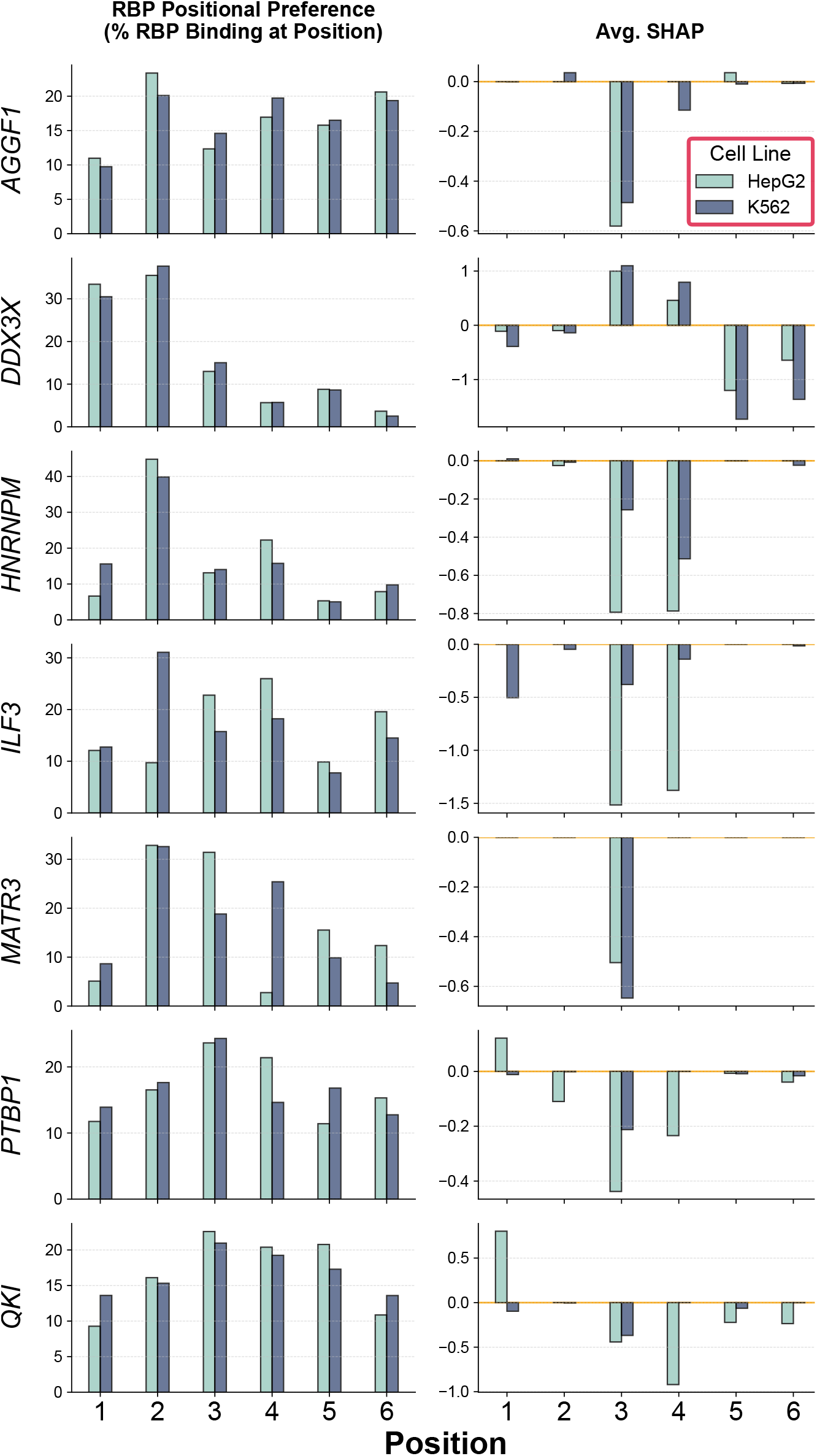
The left column calculates the percentage of an RBP’s total binding at a given position across all SIs whereas the right column indicates the Avg. SHAP at each position for that RBP.

**Figure S4.**
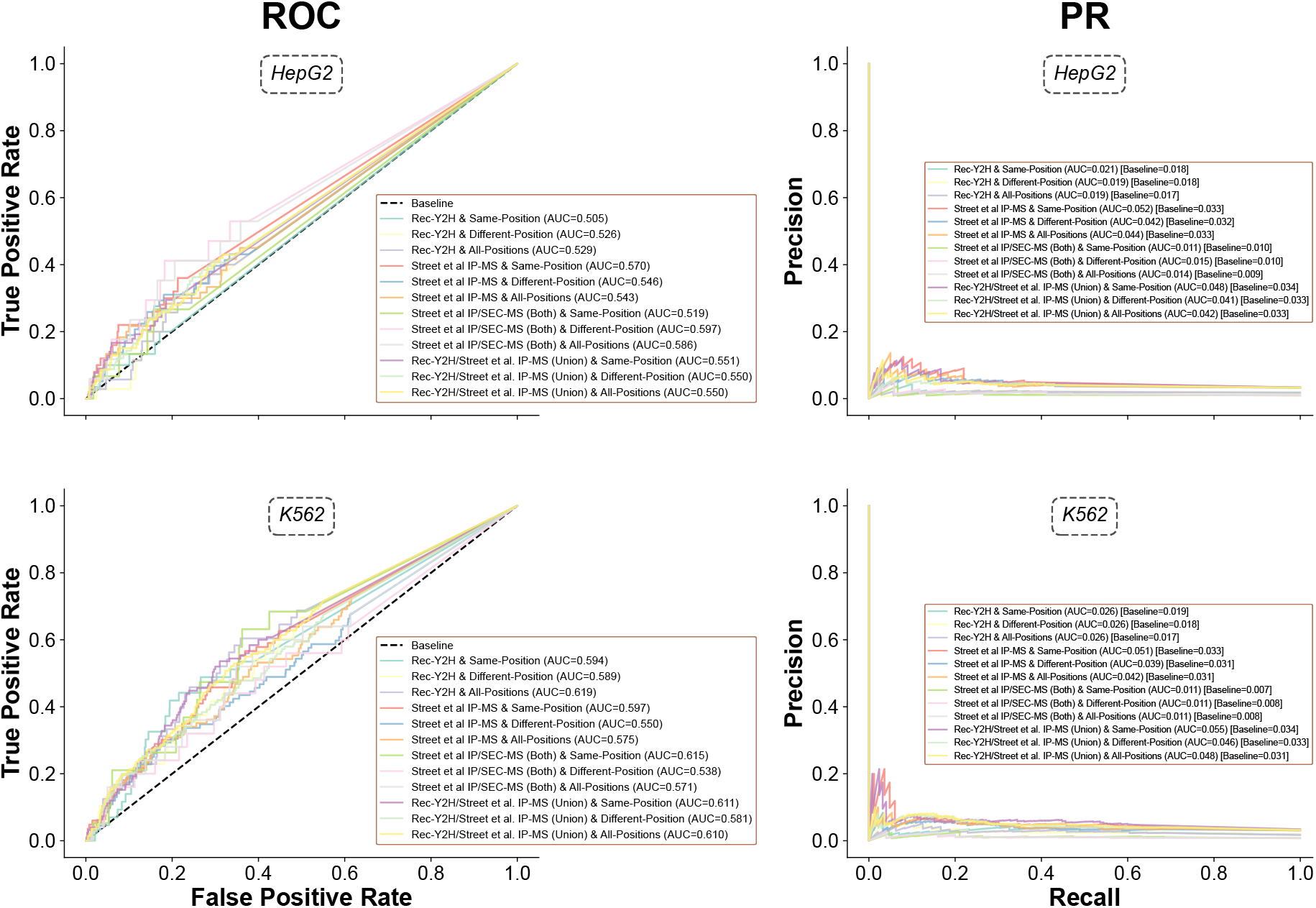
Receiver Operating Characteristic (ROC) and Precision Recall (PR) curves from using “Avg. |Interaction SHAP|” (Methods 4.12) magnitude to decipher Protein-Protein Interaction (PPI) status across multiple PPI resources (Methods 4.13) and position types in both cell lines (Methods 4.14). For each RBP pair, SHAP values were summed across positions to obtain a single summary value for each RBP pair. Please see Methods 4.14 for details on creation of ROC and PR curves. Area Under Curve (AUC) values are provided for each curve in each subplot.

**Figure S5.**
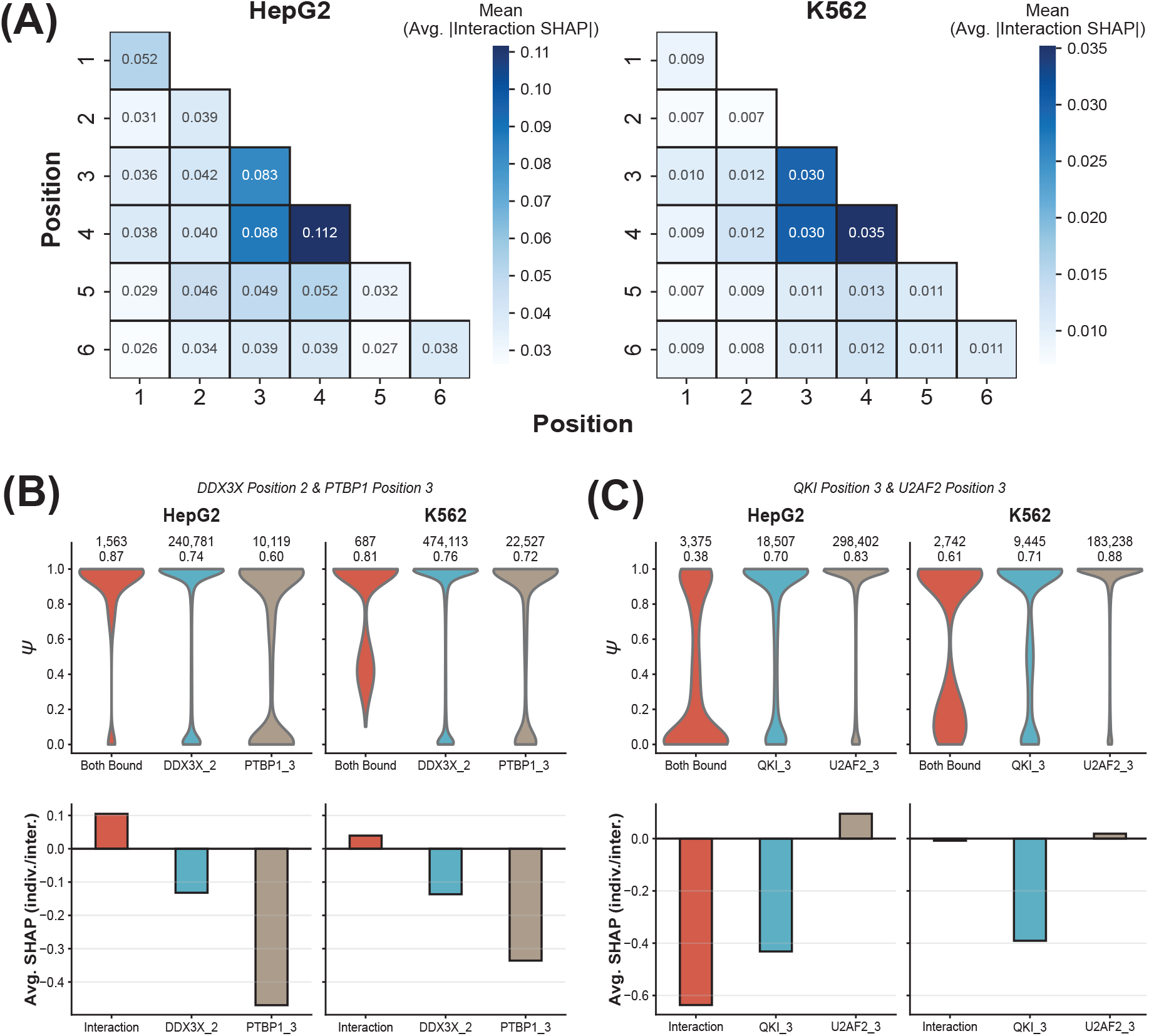
**(A)** Average of all Avg. |Interaction SHAP| values (Methods 4.12) per each position-position combination. **(B)** (Top row) *ψ* distributions for where either DDX3X is bound at position 2, PTBP1 is bound at position 3, or both. Above each violin plot is the number (top) and the average (bottom) of the *ψ* values. (Bottom row) the Avg. Interaction SHAP (left), Mean of individual-effect SHAP values (middle, right). For details, please see Methods 4.15. **(C)** Similar to **(B)** except looking at the interaction between QKI at position 3 and U2AF2 at position 3.

## 7 Supplementary Tables

**Table S1**. Matthew’s Correlation Coefficient (MCC) values for Δ*ψ* versus local SHAP using all CRISPRi RBP knockdown examples.

**Table S2**. All “Avg. SHAP”, “Avg. |SHAP|”, and “Activity” scores for each feature (an RBP at a position) in both cell lines.

**Table S3**. All “Avg. Interaction SHAP” and “Avg. |Interaction SHAP|” values for every feature-feature interaction across both cell lines along with the PPI status (using multiple PPI resources) of the RBPs involved in a given interaction.

**Table S4**. Summary statistics after testing for significant changes in actual *ψ* values when looking at feature-feature SHAP interactions where the interaction and individual effects all had to be sign concordant across both cell lines.

**Table S5**. Summary statistics after testing for significant changes in actual *ψ* values when looking at feature-feature SHAP interactions where the interaction effect had to be larger than both individual effects, with no requirement for sign concordance between cell lines.

